# Non-targeted multimodal metabolomics data from ovine rumen fluid fractions

**DOI:** 10.1101/2021.01.10.426113

**Authors:** Nikola Palevich, Paul H. Maclean, Mingshu Cao

## Abstract

From an animal production and health perspective, our understanding of the metabolites in ruminant biofluids, particularly rumen fluid across different host species is poorly understood. Metabolomics is a powerful and sensitive approach for investigating low molecular weight metabolite profiles present in rumen biofluids. It can be used to identify potential roles of metabolites in the rumen microbiome and provide and understanding of host-level regulatory mechanisms associated with animal production. The rumen is a strictly anaerobic environment enriched with a complex community of bacteria, protozoa, fungi, archaea and bacteriophages. Here, we present a metabolomic dataset generated using hydrophilic interaction liquid chromatography (HILIC) and semi-polar (C18) chromatography methods coupled to high resolution mass spectrometry (MS), collected in both positive and negative ionization modes, of ovine rumen samples that were fractionated based on molecular weight (20 kDa, 8-10 kDa and 100 Da). This study highlights the potential of HILIC and C18 chromatography combined with non-targeted mass spectrometric methods to detect the polar and semi-polar metabolite species of the ruminal fluid metabolome.

## ANNOUNCEMENT

Ruminant livestock are an important component of feeding the growing human population while also being sources of global greenhouse gas emissions. Rumen microbiota breakdown and convert plant proteins and polysaccharides from feed into energy sources but also result in methane formation that affect ruminant productivity. While rapid developments in genomics have accelerated our knowledge of rumen molecular biology (1), there has been less work focusing on the low molecular weight molecules that stem from rumen fermentation of feed and a complete absence of metabolomics studies on ovine rumen samples (2-8).

Whole rumen content samples were collected post-mortem and pooled from five sheep (Fig. 1A) grazing *ad libitum* on a ryegrass and clover pasture diet in Palmerston North, New Zealand (40°18′ S, 175°45′ E). A method was developed to acquire dialyzed rumen fluid (DRF) fractions that enrich for different sized components (Fig. 1B). DRF fractions based on three molecular weight cut-offs (MWCO) were obtained using Spectra-Por® Float-A-Lyzer® G2 dialysis systems with MWCOs of 20 kDa (Z726931, Sigma-Aldrich), 8-10 kDa (Z726605, Sigma-Aldrich) and 100 Da (Z727253, Sigma-Aldrich). Approximately 5 L of rumen contents were collected from each animal and filtered through 4 layers of cheesecloth (335 μm mesh) to account for the macro components of rumen fluid and transferred into Schott gas washing bottles fitted with Drechsel type head connections (GL 14, DURAN). To obtain each DRF fraction, replicates of each individual MWCO apparatus (*n* = 5 for each) were dialyzed against 10 mL of autoclaved phosphate buffered saline buffer overnight at 39□ in a water bath under anaerobic conditions obtained by insufflating a stream of O_2_-free CO_2_ inside the container with constant mixing. DRF samples were snap-frozen in liquid nitrogen, transferred to glass vials and stored at - 80°C until further use.

**FIG 1.**
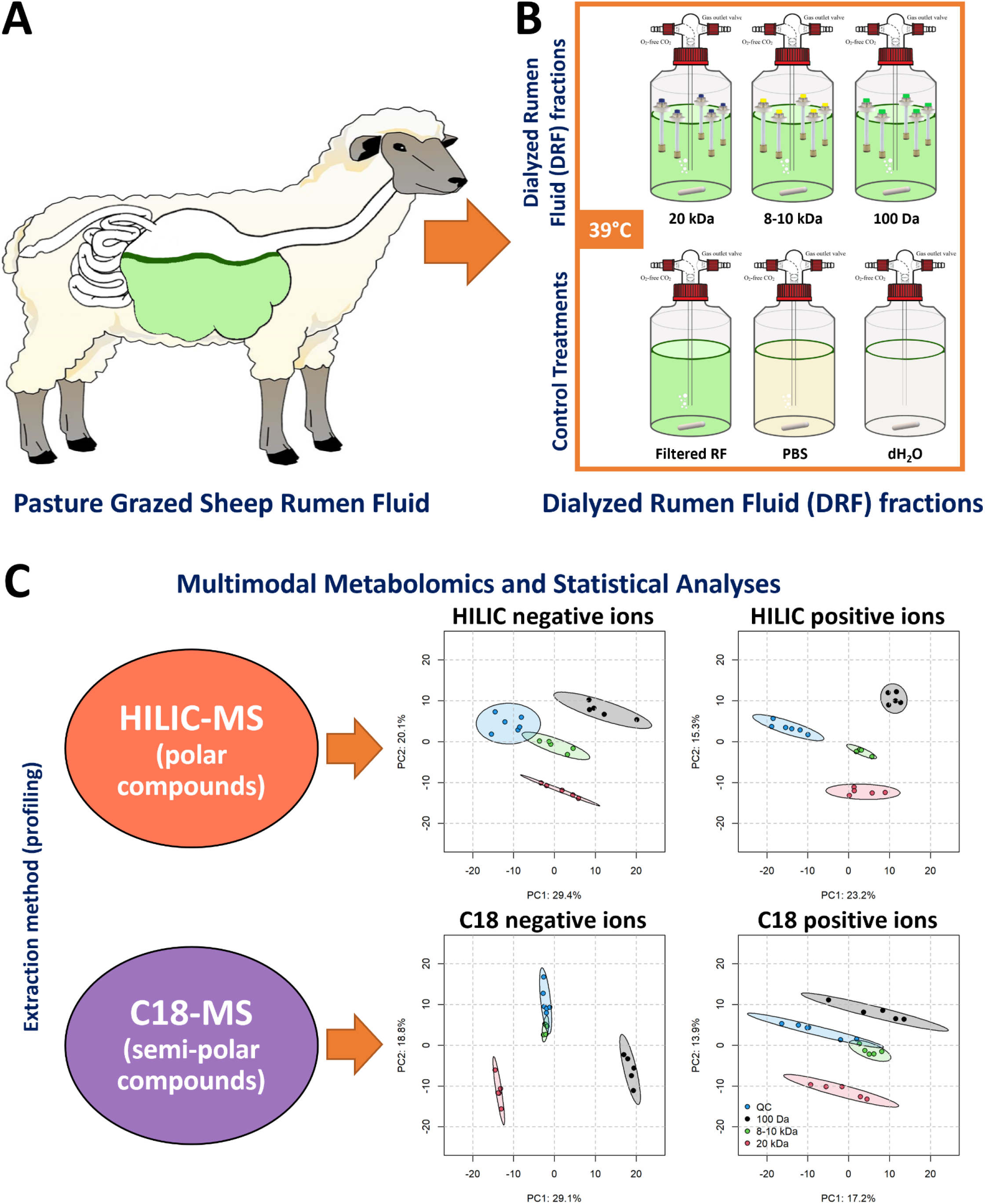
Overview of the experimental design, multimodal metabolomics workflow and statistical analysis. (**A**) New Zealand pasture-fed sheep used for this study. (**B**) Dialyzed rumen fluid (DRF) fractions were obtained under anaerobic rumen conditions (39L and CO_2_) using dialysis systems at three molecular weight cut-offs. (**C**) Schematic of a multimodal metabolomics workflow and statistical analyses used to process data integrated from multiple analytical approaches. Multigroup DRF fraction metabolic profiles across two metabolomics analyses. Principal component analysis (PCA) scores plot for the first two components of raw chromatographic peaks for C18 column negative ions (top left), C18 column positive ions (top right), HILIC column negative ions (bottom left) and HILIC column positive ions (bottom right). Each MWCO or DRF fraction was coloured to represent 20 kDa (red), 8-10 kDa (green), 100 Da (black) and QC control (blue) samples. The QC comprised a pooled aliquot of the extract of all samples. The scatter plots shown represent plots with the 2 components having the greatest variations. Percentages on axes indicate the percentage of explained variance for the respective principal components. Ellipses are 95% confidence intervals of the scores of dialyzed rumen fractions. Observations that are similar will fall close to each other in clusters. One data point represents one biological replicate. Abbreviation: HILIC; hydrophilic interaction liquid chromatography.

To comprehensively survey metabolites associated with DRF fractions we used: hydrophilic interaction liquid chromatography (HILIC) to separate polar compounds and C18 chromatography for separation of semi-polar compounds (Fig. 1C). Analyses were performed in both positive and negative electrospray ionization (ESI) mode at the resolving power setting of 25,000 with a maximum trap fill time of 100 ms using the Xcalibur v4.3 software. The LC-MS raw data files were converted to mzXML files using MSConvert function of ProteoWizard™ v3 software (9). Quality control and peak deisotoping analysis was based on our previously published procedures (10). The details of extraction procedures, chromatographic gradients, instrument settings have been previously described by Palevich, et al. (11). For each metabolomic dataset, principal component analysis (PCA) was performed on the log_10_ intensities for raw chromatographic peaks to assess similarity and separation of DRF fractions. The *mixomics* package v6.16.3 in R v4.1.1 was used to perform the PCA and generate plots.

Overall, according to the PCA scores plots (Fig. 1C), each of the three DRF fractions had considerably different profiles for both types of metabolomic analyses and regardless of ionization mode. The three DRF fractions separated completely either on the first or second component or a combination thereof within the 95% confidence interval ellipse. The presented untargeted metabolomics data provides a detailed snapshot of the ovine ruminal fluid metabolome that can be used as a reference for future studies of the rumen metabolome or as a comparator for other ruminant species.

### Data availability

The datasets presented in this study and supporting the conclusions of this article have been made available in the MetaboLights database **[MTBLS1717]** online repository.

## ACKNOWLEDGEMENTS

This work was funded by the AgResearch Ltd. Strategic Science Investment Fund (SSIF, Grant No. C10X1702) to NP. This work was also supported in part by the Agricultural and Marketing Research and Development Trust (AGMARDT) Postdoctoral Fellowship Programme (Grant No. P17001) to NP. Special thanks to Linley Schofield in our Rumen Microbiology lab for providing the dialysis equipment, her guidance and advice on using the apparatus. We thank Hailey Gillespie and Trevor Holloway who were always willing to collaborate and especially for their timely assistance with the collection of rumen fluid. We also wish to acknowledge Arvind Subbaraj for assistance with the LCMS analysis for this study, and Alastair Ross and Karl Fraser for their feedback on early versions of this manuscript.

